# UBR4 regulates a MetAP2-dependent Arg/N-degron pathway

**DOI:** 10.1101/2024.10.03.616566

**Authors:** Evan J. Morrison, Emma Horton, Olivia S. Rissland

## Abstract

The open reading frame does more than merely encode a linear peptide sequence; it is a reservoir of regulatory information. Here, as part of investigations into how the N-terminal amino acids regulate translation, we serendipitously uncovered a new N-degron that revealed an additional layer of regulation in these pathways. Using reporter assays, we discovered that peptides bearing position 3 arginine or lysine residues at the N-terminus were rapidly degraded in mammalian cells. We found this pathway requires MetAP2, which co-translationally cleaves the N-terminal methionine preceding second position threonine and valines to initiate protein decay. We used CRISPR-Cas9 to knockout key N-recognins and found that these N-degrons are exclusively targeted by the E3 ligase UBR4, but not by UBR1 or UBR2. Together, our results characterize a new N-degron pathway that reveals a unique role for MetAP2 and UBR4 in mediating protein decay.

**SIGNIFICANCE:** The Arg/N-degron pathway targets position 1 or 2 N-terminal Lys and Arg residues via UBR Box E3 ligases to trigger protein decay. Here we show that UBR4 can specifically recognize position 3 Lys and Arg N-termini upon methionine removal by the methionine amino peptidase MetAP2. Accordingly, proteins that bear N-terminal residues that are processed by MetAP1 are unaffected by the loss or inhibition of MetAP2. Using a combination of reporter assays, and bioinformatic approaches were identified endogenous proteins whose N-termini are recognized by this MetAP2-dependent Arg/N-degron pathway. Thus, our results expand the number of Arg/N-degron substrates and describe a new mechanism through which they are targeted.

## INTRODUCTION

N-degrons are sequence elements at the very N-terminus of proteins that trigger protein decay in all known eukaryotes (1). For example, peptides bearing N-terminal lysines, arginines, or bulky hydrophobic residues are targeted by UBR E3 ligases through the Arg/N-degron pathway, while N-terminal prolines are targeted by the GID/CTLH E3 ligases via the pro/N-degron pathway (1–3). Even though the importance of amino residues at position 1 or 2 for N-degrons was discovered nearly four decades ago (4), a complete working model has yet to emerge due to the multiplicity of factors that influence their accessibility and processing.

N-degrons with non-methionine residues at position 1 must be generated through proteolytic cleavage events (1). For instance, caspases cleave peptides upstream of glycine residues to produce Gly/N-degrons (5). Other N-degrons are generated through the co-translational removal of the initiator methionine by methionine amino peptidases (3, 6–8). Adding even more complexity, recent studies show that some peptides require several successive proteolytic events such as methionine removal, followed by dipeptidyl peptidase cleavage to expose Arg/N-degron motifs (9). Additionally, co-translational acetylation of the N-terminus either before or after methionine processing can both activate and protect substrates from N-degron pathways (6, 10, 11). Similarly, E3 ligases have additional sequence specificity conferred by residues downstream of N-terminal amino acid (5, 12, 13). The dizzying combinations of regulatory constraints has made it difficult to find, characterize, and predict endogenous N-degron substrates and pathways.

Given the importance of co-translational removal of the initiator methionine, the substrate specificity of MetAPs is a critical aspect to N-degron activity. In eukaryotes, methionine removal is regulated by two distinct methionine amino peptidases, MetAP1 and MetAP2 (14). Generally, both MetAPs can cleave the methionine from peptides when the second position is glycine or cysteine, while MetAP1 exclusively targets peptides with a second position alanine, serine, or proline, and MetAP2 cleaves methionine from peptides with a position 2 threonine or valine (14–16). While removal of methionine from MetAP1 substrates is thought to be highly efficient, methionine processing by MetAP2 can be variable (14, 15). Moreover, the removal of the methionine from proteins with position 2 alanine, serine, threonine, and valine is thought to have a minimal effect on protein stability, as these are known as stabilizing amino acids (5). Thus, there is little evidence to date that MetAP2 plays a strong role in N-degron pathways (5, 7). In contrast, MetAP1 plays a critical and near exclusive role in removing the methionine from Pro/N-degrons, Ac/N-degrons, and the recently discovered DPP-dependent Arg/N-degrons, as well as GASTC/N-degrons (1, 3, 9, 17, 18). Additionally, in some rare instances, it has been shown in yeast that MetAP1 (Map1 in yeast) can remove the methionine from asparagine and glutamine containing peptides to create tertiary Arg/N-degrons (7, 8).

Another important aspect of N-degrons is their recognition by corresponding ubiquitin E3 ligases, known as N-recognins. In humans, Arg/N-degrons are targeted by the UBR E3 ligases, UBR1, UBR2, UBR4, and UBR5 (1, 2, 19). UBR E3 ligases contain a UBR Box domain, which is used to target so-called type-1 Arg/N-degrons that bear positive N-terminal residues (12, 19–21). High throughput studies of UBR1, UBR2, and UBR4 have shown that UBR4 targets a wider range of substrates such as type-1, position 2 residues (MK and MR) (5). In line with these observations, recent structural evidence points to a divergence in the UBR box that leads to different substrate recognition by UBR4 from that of UBR1 and UBR2 (21, 22). UBR4 has also attracted interest for its role in brain and cardiovascular development as well as its role in recognizing and responding to mitochondrial stress through internal basic degron sequences as part of the SIFI complex (which also contains potassium channel modulatory factor, KCMF1 (23, 24)). Despite work from multiple labs, it is unclear whether KCMF1 is required for UBR4 to recognize type-1 Arg/N-degrons as has been shown for type-2 degrons, as well as internal degrons (5, 11, 19, 25, 26).

Here, we were interested in understanding how N-terminal regions impacted gene expression, and serendipitously discovered an unanticipated Arg/N-degron pathway. We found that a position 3 lysine or arginine that follows a MetAP2 cleavage site triggers robust decay of a luciferase reporter. Notably, removal of the methionine is necessary but not sufficient to induce decay, as reporters with MetAP1-targeted position 2 amino acids were stable. Using a combination of genetic tools, we found that UBR4, but not UBR1 nor UBR2, participates in degron recognition, expanding the Arg/N-degron pathway. Our results indicate that the destabilizing effect of position 3 lysines and arginines upon methionine removal is widespread throughout the human proteome and identify endogenous N-termini that are sufficient to activate the MetAP2-dependent Arg/N-degron pathway. Crucially, the stability of these N-degrons is modulated by the position 2 residue, as well as downstream residues, likely due to substrate specificity for a variety of enzymes involved in this pathway, including MetAP2 and UBR4. Our results expand our understanding N-degrons pathways and underscore how much remains to be discovered about the complex regulatory logic of N-terminal sequences.

## RESULTS

### An N-terminal KIH motif is destabilizing

Previous research in *E. coli* demonstrated that insertion of amino acids at positions 3-5 of exogenous reporter genes, like eGFP, can substantially alter reporter expression by controlling translation elongation processivity (27). For instance, the addition of tripeptide motif of lysine– isoleucine–histidine (KIH) at this position increased expression by an order of magnitude compared to unmodified eGFP (27). We initially wondered if the same result would hold in human cells and opted to use dual luciferase assays to test the generality of the observation. Here, we inserted the KIH tripeptide motif at positions 3–5 of *Renilla* luciferase (Figure 1A) and expressed the KIH or unmodified *Renilla* luciferase in HEK293T cells together with a firefly luciferase control. Strikingly, rather than having higher expression (as occurred in *E. coli*), the KIH-containing reporter exhibited ∼20-fold reduced expression compared to control *Renilla* luciferase (Figure 1B; *p* < 10^−5^). Similarly, when we changed the optimality of codons in the KIH insertion from AAG-ATC-CAC to AAA-ATA-CAT, the luciferase expression was equally reduced demonstrating that the loss of expression was primarily driven by the identity of the amino acids rather than the codons (Supplemental Figure 1A). We obtained the same result using western blotting, indicating that the decrease in luciferase activity was due to a decrease in expression rather than a decrease in enzymatic activity (Figure 1C). To test the extent to which this observation was cell-type specific, we transfected these reporters into three distinct human cells lines, U2OS, RPE1, and K562, along with two non-human mammalian cells lines, NIH/3T3 (a mouse embryonic fibroblast cell line), and CHO-K1 (a Chinese hamster ovarian cell line); and the KIH insertion strongly decreased luciferase expression across all cell lines (Supplemental Figure 1B, C). These results suggest that the KIH motif broadly represses expression across mammalian cell types.

**Figure 1.**
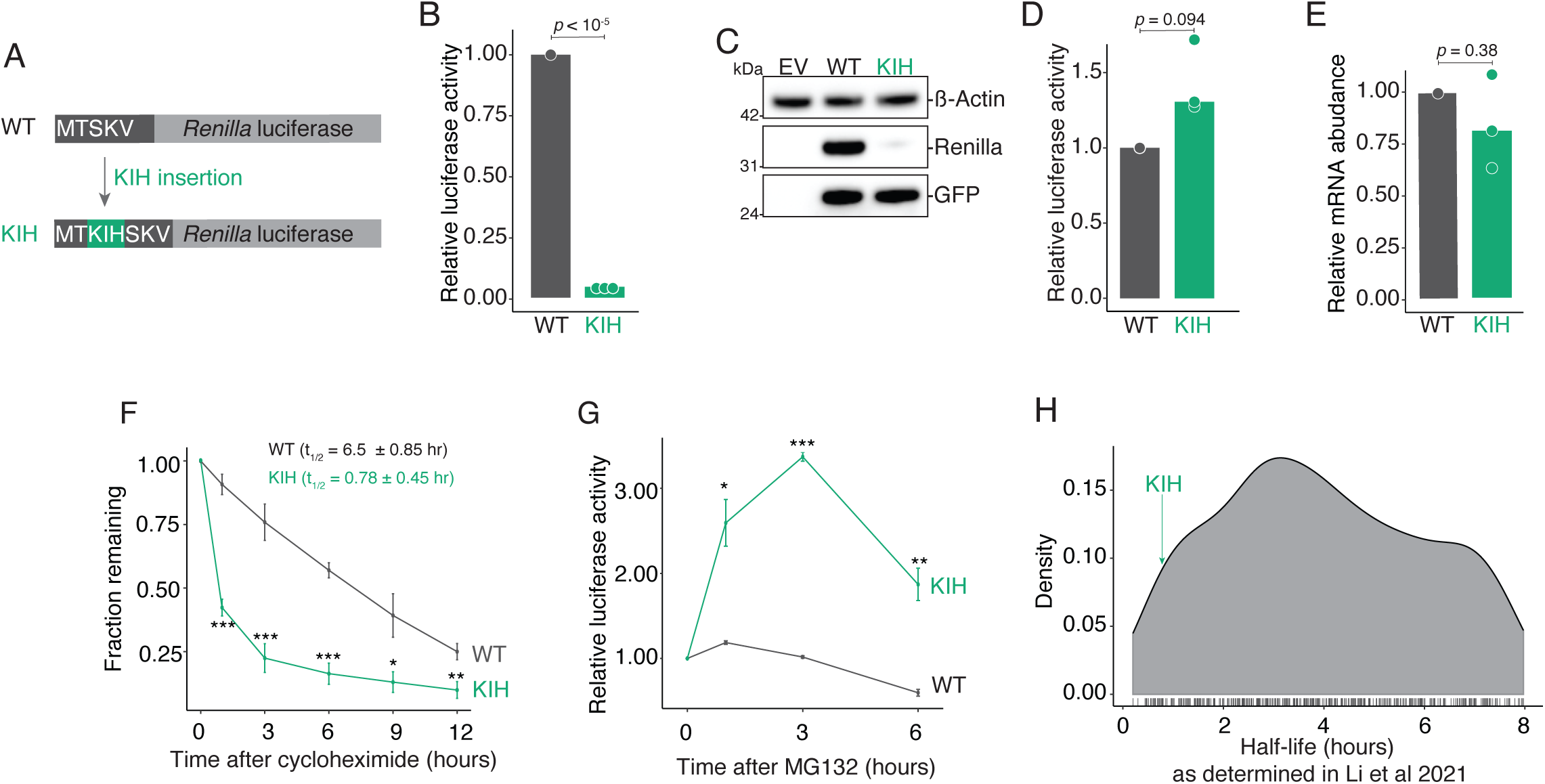
An N-terminal KIH motif is destabilizing. (A) Schematic of reporter constructs. We inserted KIH codons (AAG ATC CAC) into positions 3-5 (shown in green) of a codon optimized *Renilla* luciferase reporter. An unmodified version was used as the “wild-type” comparator. (B) KIH *Renilla* luciferase has decreased activity. HEK293T cells were transfected with either wild-type or KIH *Renilla* luciferase variants, and co-transfected with firefly luciferase as a control. *Renilla* luciferase activity was quantified relative to the firefly control, and *Renilla* expression was calculated relative to the wild-type version. The bar represents the median activity, and each dot represents one biological replicate (composed of three technical replicates). P-value was calculated using the Mann-Whitney U test. (C) KIH *Renilla* has decreased protein expression. HEK293T cells were transfected with empty vector (EV), WT or KIH *Renilla* luciferase, and co-transfected with GFP as a transfection control. Lysates were made and run by gel electrophoresis, and membranes probed for *Renilla* luciferase, GFP, and β-actin. (D) The KIH motif does not reduce expression in *in vitro* systems. Reporters were *in vitro* transcribed and incubated in rabbit reticulocyte lysate. *Renilla* luciferase activity was quantified as in (B), and P-value was calculated using Student’s t-test. (E) RNA levels do not explain the difference between wild-type and KIH expression levels. Transfection as in (B), *Renilla* luciferase RNA levels were quantified with RT-qPCR normalized to firefly luciferase levels, and P-value was calculated using Student’s t-test. (F) KIH luciferase is rapidly degraded. Cells were transfected as in (B) for 24 hours and then treated with 80 μg/mL of cycloheximide (CHX) or DMSO for the indicated time points. Luciferase assays were used to determine the amount of *Renilla* luciferase, and the fraction remaining was normalized to the 0-hour time point for each luciferase variant. Student’s t-test was used to determine the statistical significance between luciferase variants for each time point. A * indicates *p* < 0.05; ** *p* < 0.01; *** *p* < 10^−3^. (G) KIH luciferase expression is rescued by proteasome inhibition. As in (F) except cells were treated with 10 μM MG132 at the indicated time points. (H) The KIH luciferase variant is less stable than 99% of endogenous proteins in HEK293T cells. The bottom 5% of short-lived proteins were plotted according to their half-lives. The half-life of KIH *Renilla* luciferase is indicated by the green arrow. Data obtained from Li *et al.* 2021.

We next wondered what layer of gene regulation was responsible for the decrease in KIH expression. We first considered that the motif might mediate decreased early translational elongation processivity, in contrast to the scenario in *E. coli*; however, when the variants were expressed in a rabbit reticulocyte *in vitro* translation system, the KIH version now had slightly *higher* expression (1.31-fold, p = 0.09, Figure 1D). This result argued against a strong translational effect that we would have predicted to be recapitulated in this system. Another possibility was that the addition of the KIH codons destabilized the reporter mRNA. However, we observed no significant difference in mRNA abundance as determined by qRT-PCR between our control and KIH variants (*p* = 0.38; Figure 1E).

Finally, we explored whether the KIH tripeptide motif resulted in faster degradation of the *Renilla* reporter. We initially had excluded this possibility because, at the time, no position 3 amino acid degron had been reported, and the position 2 amino acid in our reporter (threonine) does not commonly act as an N-degron. To test this hypothesis, we measured the half-lives of our WT and KIH reporters using cycloheximide chase assays. While the half-life of the wild-type *Renilla* luciferase was 6.5 ± 0.85 hours, the half-life of KIH version was significantly reduced to 0.78 ± 0.45 hours (*p* < 10^−3^; Figure 1F). To complement these experiments, we examined the impact of the proteasomal inhibitor MG132 on the two luciferase variants. Consistent with the measured half-lives, the expression of KIH version increased rapidly after the addition of MG132 while wild-type variant did not (Figure 1G).

Recently, it has been shown that ∼95% of endogenous proteins have a half-life greater than 8 hours (28). As such, we note that our KIH luciferase variant is exceptionally short-lived. Indeed, comparing the stability of the KIH luciferase with the stability of the shortest-lived proteins in HEK293T cells showed that this luciferase variant was less stable than 95% of these proteins (Figure 1H) (28). Taken together, these results demonstrate that, KIH at positions 3–5 is a potent degron in human cells.

### KIH and other related tripeptide motifs act as N-degrons

We next wondered whether the impact of the KIH motif was position dependent and created reporters where its position was shifted across the N-terminus one amino acid at a time, from position 3-5 to 4-6, 5-7, etc. (Figure 2A, B). As an internal control, firefly luciferase was expressed from the same transcript using an EMCV IRES downstream of the *Renilla* luciferase. Although KIH at other positions did modestly reduce expression by ∼2-fold, only the motif at position 3-5 resulted in the order of magnitude loss we originally observed. As a complimentary approach, we also monitored the decay rates of the reporter proteins, and, consistent with the steady state measurements, only KIH at position 3-5 significantly destabilized the *Renilla* reporter (Figure 2C). These results demonstrate that KIH, despite not following any known rules of N-degrons, is a position-dependent N-terminal degron.

**Figure 2.**
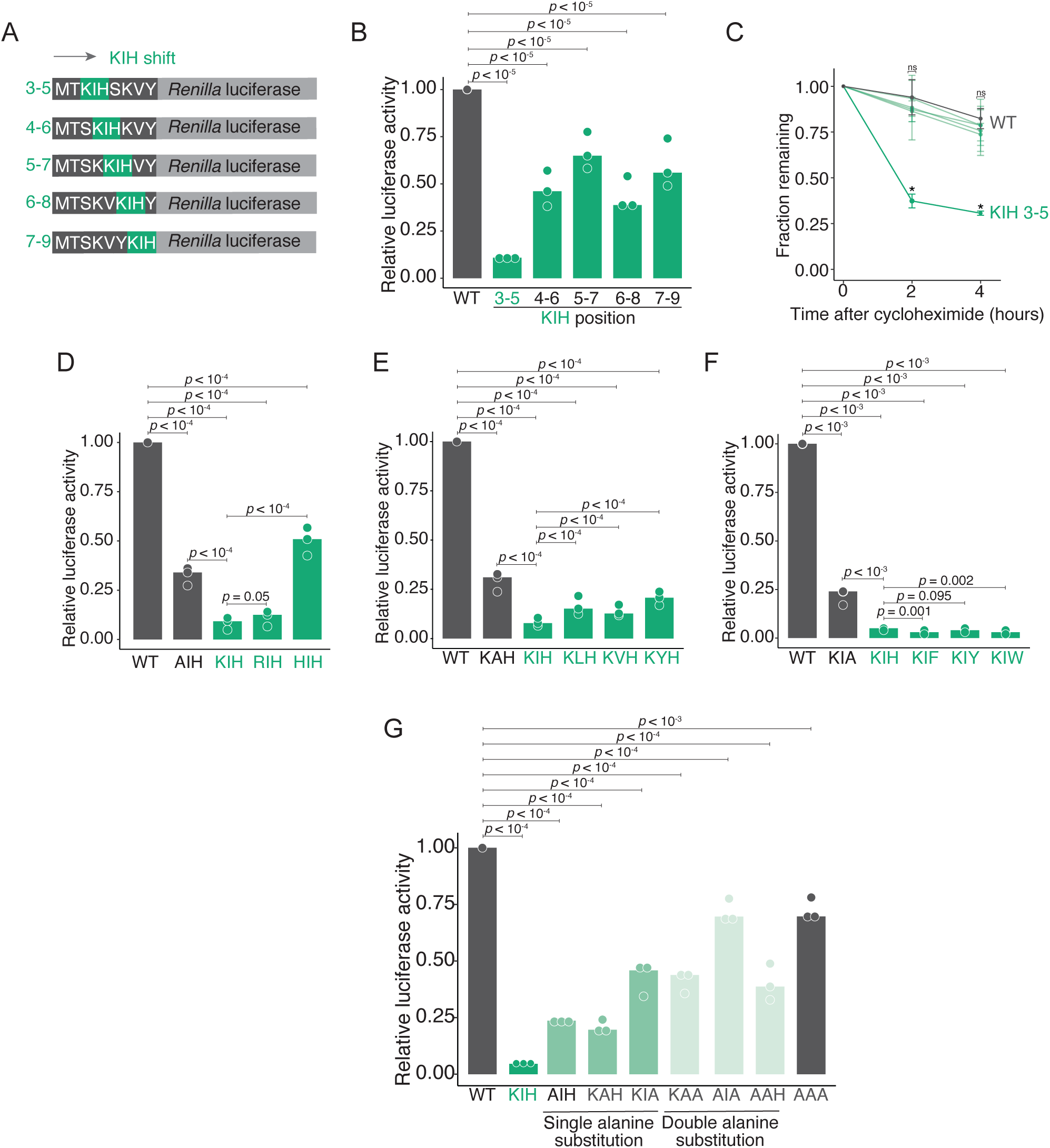
KIH and other related tripeptide motifs act as N-degrons. (A) Schematic of reporter constructs. We shifted KIH codons (AAG ATC CAC) into positions 4-6, 5-7, 6-8, and 7-9 (shown in green) of a codon-optimized *Renilla* luciferase reporter. Reporters also contained a firefly luciferase expressed from the same transcript with the minimal EMCV IRES. (B) KIH-mediated loss of luciferase expression is position dependent. HEK293T cells were transfected with either wild-type or KIH *Renilla* luciferase variants in (A). *Renilla* luciferase activity was quantified relative to the firefly control, and *Renilla* expression was calculated relative to the wild-type version. The bar represents the median activity, and each dot represents one biological replicate (composed of three technical replicates). P-value was calculated using the Mann-Whitney U test. (C) KIH mediated decay is position dependent. Cells were transfected as in (B) for 24 hours and then treated with 80 μg/mL of cycloheximide (CHX) or DMSO for the indicated time points. Luciferase assays were used to determine the amount of *Renilla* luciferase, and the fraction remaining was normalized to the 0-hour time point for each luciferase variant. Student’s t-test was used to determine the statistical significance between luciferase variants for each time point. A ns indicates *p* > 0.05; * indicates *p* < 0.05. (D) A lysine or arginine at position 3 decreases *Renilla* luciferase protein levels. As in (B) except with the indicated reporters. (E) Bulky hydrophobic residues at position 4 reduce luciferase expression. As in (B) except with the indicated reporters. (F) Aromatic residues at position 5 reduce gene expression. As in (B) except with the indicated reporters. (G) Alanine mutations are sufficient to increase luciferase expression. As in (B) except with the indicated reporters.

We next turned to understanding the rules of this novel degron. Position 2 lysine is notable in acting as a robust N-degron, but it can also serve as the site for ubiquitylation (1, 5). To separate these potential roles, we swapped lysine for other positively charged amino acids (arginine and histidine) or alanine, as a control, and measured the impact via dual luciferase assays as before. While all tripeptide insertions decreased *Renilla* luciferase activity relative to the wild-type version, only RIH showed an effect similar to KIH, suggesting that at a positive charge at position 3 is a prominent feature of the degron (8-fold, *p* < 10^−4^; Figure 2C) and that the ubiquitylation at this position is not required. We then swapped the position 4 isoleucine to similar hydrophobic residues. All versions resulted in a strong decrease in expression, although an alanine at this position (as part of KAH) was unable to mediate as robust of an impact; these results indicate bulky hydrophobic residues at position 4 also enhances the decay (Figure 2E). Next, we swapped histidine at position 5 for other aromatic residues (phenylalanine, tyrosine, and tryptophan) as well as alanine, as a control (Figure 2F). Interestingly, despite the charge difference between histidine and Phe/Tyr/Trp, all four were able to function as degron sequences, with KIF and KIW having an even larger destabilizing effect (Figure 2F). Lastly, we systematically replaced positions within the motif with alanine; as an example, we examined the impact of the single replacement series (AIH, KAH, and KIA) on *Renilla* luciferase protein expression (Figure 2G). Replacing any of the three positions with alanine weakened the destabilizing effect, and the double replacements further reduced the impact of the motif. Thus, no single amino acid or pair of amino acids within the tripeptide motif is sufficient for the N-degron. Together, these results indicate that KIH functions as an N-terminal degron that is part of a larger family of degrons described by a tripeptide of [K/R]-[semi-bulky hydrophobic]-[aromatic amino acids].

### Removal of the initiator methionine by MetAP2 is required for the KIH degron

To our knowledge, studies of N-degron pathways to date have primarily focused on the role of the first or second amino acid in controlling decay (1). Given that KIH triggered decay when it was at positions 3-5 of *Renilla* luciferase, it was unclear which N-degron pathway mediated this effect. Nonetheless, a common theme for N-degrons is removal of the initiator methionine, and we reasoned that the same might be the case here. We made use of ubiquitin-fusion reporters, whereby we inserted a ubiquitin immediately upstream of *Renilla* luciferase in our wild-type or KIH reporters (Figure 3A). Upon translation of the ubiquitin-fusion reporter, deubiquitinases rapidly co-translationally cleave the ubiquitin exposing the programmed N-terminus (29, 30). As before, these reporters also contained an internal control firefly luciferase driven by an EMCV IRES. Previous studies have shown that the methionine at position 1 after ubiquitin cleavage is not removed by MetAPs (5, 30). Strikingly, when position 1 methionine was included with the KIH variant (giving an N-terminus of MTKIH), expression levels were 80% that of the wild-type version. In contrast, for the reporter with the TKIH N-terminus (i.e., where the “position 1” methionine has been removed), we again observed a ∼10-fold decrease in expression. This result indicates that removal of the methionine is necessary for rapid decay mediated by the KIH degron.

Methionine removal is controlled by the radius of gyration of the second amino acid (14, 16). Amino acids with a small radius of gyration (G, A, S, C, T, and V) are cleaved by methionine amino peptidases (MetAPs), while amino acids with large radii of gyration are not cleaved (5, 14, 16). Given that the removal of the initiator methionine was necessary for KIH luciferase decay, and our original reporter included threonine at position 2, we hypothesized that the second amino acid would affect the potency of the degron where smaller residues at the second position would result in similarly low levels of expression, while larger residues would inhibit decay resulting in higher levels of expression (Figure 3B). As expected, based on a lack of any N-degron in the wild-type version, mutation of the second-position threonine had a minimal effect on luciferase levels in this context. Surprisingly, and contrary to our hypothesis, mutation of threonine to smaller residues (A and S) also increased levels of the KIH variant. This result was exceptionally notable for the T2S KIH mutation, which had ∼13-fold higher (*p* < 10^−4^) expression than the original KIH version. On the other hand, replacing the second-position threonine with larger residues increased KIH luciferase levels as predicted (Figure 3B, see Q, D, E). Moreover, consistent with our hypothesis, swapping in valine at position 2 also resulted in highly reduced levels of the KIH luciferase, although it also substantially repressed WT levels (*p* < 10^−3^; Figure 3C).

**Figure 3.**
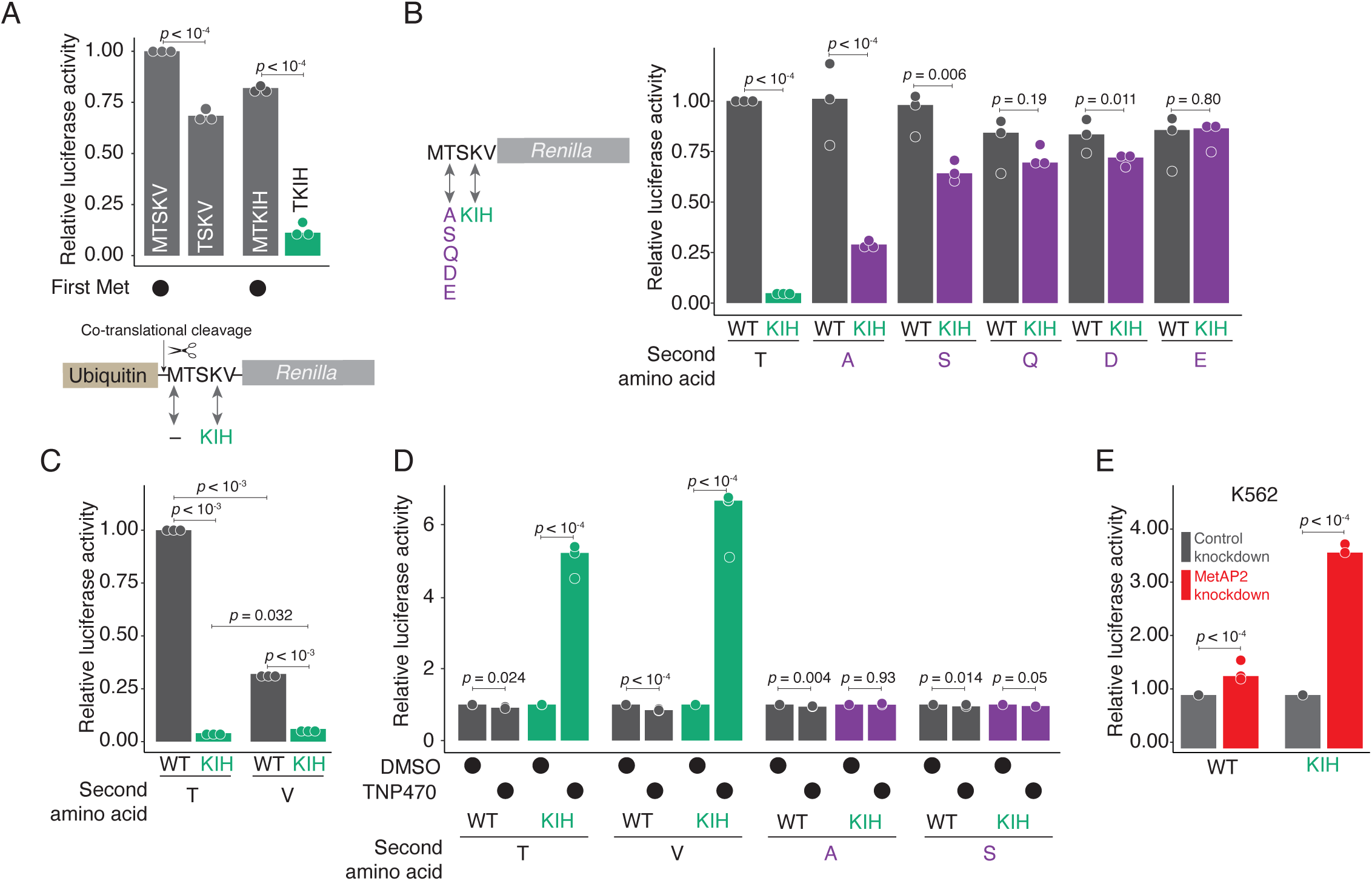
Removal of the initiator methionine by MetAP2 is required for the KIH degron. (A) Removal of the initiator methionine is required for KIH mediated decay. Schematic of reporter constructs shown below the plot. A ubiquitin moiety was placed upstream of *Renilla* luciferase variants either containing or lacking a position 1 methionine. The reporters were transfected into HEK293T cells, and *Renilla* luciferase activity was quantified relative to the firefly control. Variant *Renilla* expression was calculated relative to the wild-type version (MTSKV). The bar represents the median activity, and each dot represents one biological replicate (composed of three technical replicates). P-value was calculated using the Mann-Whitney U test. (B) The position 2 amino acid controls KIH luciferase expression. Schematic of reporter mutations are shown to the left of the plot. The second amino acid (T in the WT version) were mutated to alanine (A), serine (S), glutamine (Q), aspartic acid (D), and glutamic acid (E) in both WT and KIH luciferase variants. Cells were transfected as in (A) and *Renilla* expression for each variant was calculated relative to the WT (T) version. (C) A second position valine reduces *KIH Renilla* luciferase expression. As in (B) except with a threonine to valine mutation (T2V). (D) Removal of the N-terminal methionine by MetAP2 regulates KIH expression. HEK293T cells were transfected as in (B) with the indicated reporters. After 24 hours, cells were then supplemented with DMSO or 50 nM of TNP-470. Twenty fours after the addition of the inhibitor, relative *Renilla* luciferase activity was quantified as in (B) and luciferase expression was normalized to DMSO treated cells. Depletion of MetAP2 stabilizes KIH luciferase variants. Zim3-dCas9 containing K562 cells were transduced with either non-targeting (NT) or MetAP2 targeting gRNAs. Cells were transfected with the indicated reporters, and relative luciferase activity was quantified as in (A) and *Renilla* luciferase expression was normalized to the cells transduced with non-targeting (NT) gRNAs.

One notable difference between threonine and valine at position 2 versus alanine and serine is the identity of the methionine aminopeptidase that removes the initiator methionine (14–16). That is, MetAP2 mediates the removal of the N-terminal methionine preceding threonine and valine, while MetAP1 does so for alanine and serine (14–16). We therefore asked if MetAP2 regulated the expression of KIH luciferase with threonine and valine at position 2. To do so, we transfected cells with either wild-type or KIH *Renilla* variants with different second-position amino acids (namely, T, V, A, and S), and then added TNP-470, a cell-permeable inhibitor of the methionine aminopeptidase activity of MetAP2, or DMSO as a control (15, 31). Inhibition of MetAP2 significantly stabilized KIH luciferase expression only in the presence of a second position threonine or valine (*p* < 10^−4^), but had minimal effect on those reporters with alanine or serine at position 2 (Figure 3D). As an alternative approach, we depleted MetAP2 in K562 cells with the Zim3-dCas9 CRISPR-interference system (32, 33) (CRISPRi; Supplemental Figure 2A). Consistent with our results with TNP-470, knockdown of MetAP2 stabilized the KIH *Renilla* luciferase but not the wild-type one (Figure 3E). These results show that removal of the methionine by MetAP2 is necessary to trigger KIH decay and thus that the impact of the degron is modulated by the amino acid identity at position 2.

### UBR4 targets position 3 lysines and arginines after methionine cleavage

Having established that the removal of the methionine was necessary for the activity of the KIH degron, we recognized that the processed N-terminus now resembled known type-1 primary Arg/N-degrons that were recently characterized by a lysine at position 2 preceded by a methionine (MK) (5). Consistent with such an idea, when we examine the impact of KIH in the context of ubiquitin fusions, we found that constructs with N-termini of TKIH, VKIH, MKIH had similar expression levels in HEK293T cells (p = 0.18 to 0.62), while that of KIH had even further reduced levels (*p* < 10^−3^, Figure 4A).

**Figure 4.**
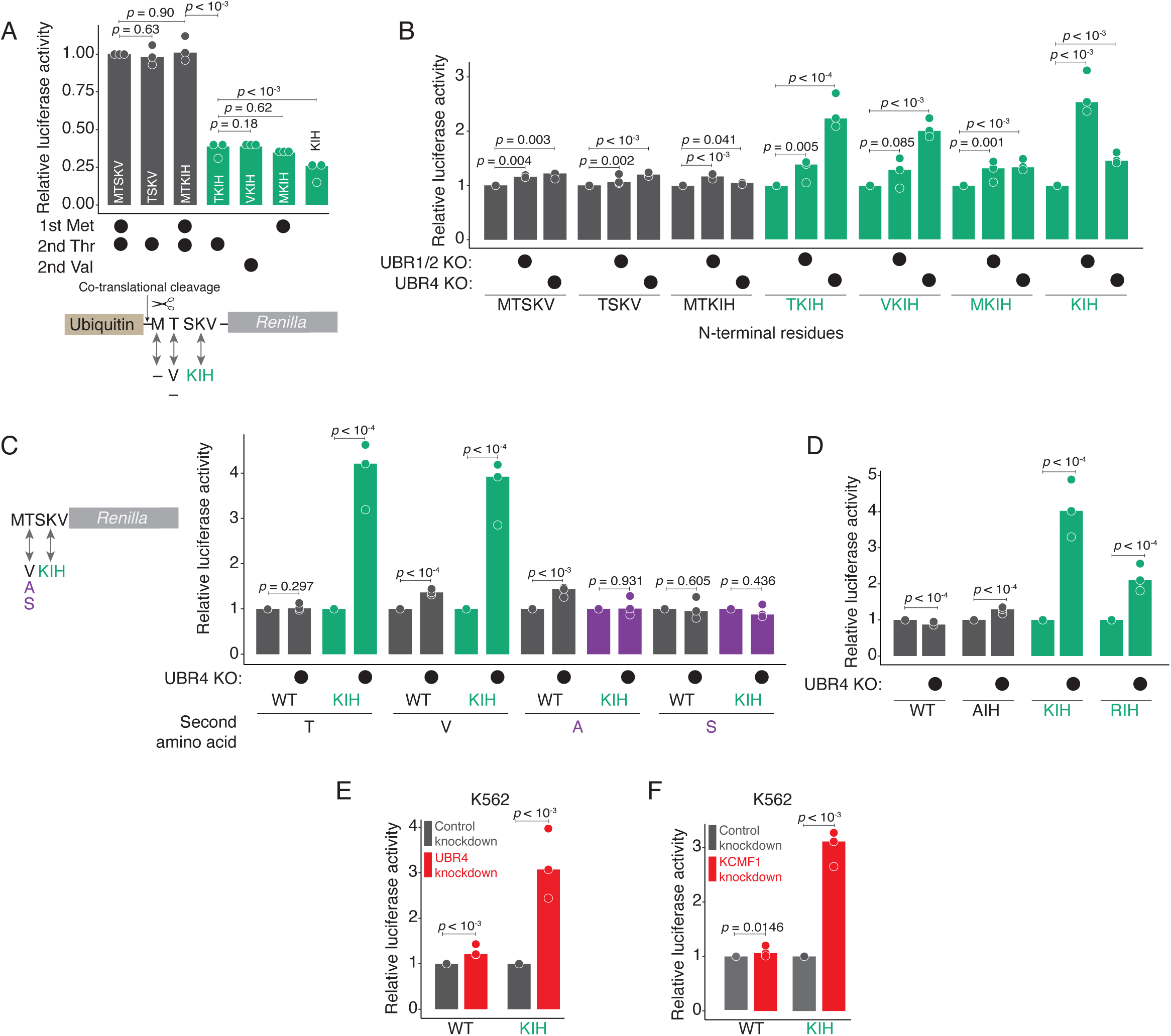
UBR4 targets position 3 lysines and arginines after methionine cleavage. (A) Position 3 lysines decrease expression as much as position 2 lysines. Schematic of reporter constructs indicated below the plot. Ubiquitin moieties were placed upstream of luciferase variants with the following N-termini: MTSKV, TSKV, MTKIH, TKIH, VKIH, MKIH, and KIH. Reporters were transfected into HEK293T cells, and relative *Renilla* luciferase activity was determined by dual luciferase assays. *Renilla* expression was calculated relative to the WT *Renilla* luciferase variant (MTSKV). The bar represents the median activity, and each dot represents one biological replicate (composed of three technical replicates). P-value was calculated using the Mann-Whitney U test. (B) UBR4 targets position 3 lysine residues that lack an initiator methionine. Reporters were transfected in parental, UBR1/2 KO or UBR4 KO cells as in (A). *Renilla* expression for each luciferase variant was calculated relative to the expression of that variant in parental cells. (C) UBR4 recognizes MetAP2 processed substrates. Schematic of reporter constructs shown below the plot. Parental or UBR4 KO cells were transfected with the indicated luciferase variants as in (B). (D) UBR4 targets position 3 arginines. As in (C) except with the indicated constructs. (E) UBR4 targets KIH luciferase across cell types. Zim3-dCas9 containing K562 cells were transduced with either non-targeting (NT) or UBR4 targeting gRNAs. Cells were transfected with the indicated reporters and *Renilla* luciferase activity was quantified as in (A) and *Renilla* luciferase expression was normalized to the cells transduced with non-targeting (NT) gRNAs. (F) The UBR4-KCMF1 complex is necessary to degrade KIH luciferase. As in (E) except with gRNAs that target KCMF1.

Primary type-1 Arg/N-degrons are targeted by the UBR Box E3 ligases (also known as N-recognins) UBR1, UBR2, UBR4, and UBR5 (1, 2, 19). Biophysical and structural studies have shown that the UBR domain of UBR1 and UBR2 has a strong affinity for tetrapeptides bearing a position 1 lysine or arginine (12, 20, 21), including the tetrapeptide KIFS, which we unknowingly tested as a motif at positions 3-6 previously (Figure 2F). The UBR box domain of UBR4 and UBR5 are also capable of recognizing these sequences, with UBR4 having a higher affinity than UBR1 or UBR2 for lysine and arginine at position 2 following the initiator methionine (5, 19). To investigate the role of UBR family of E3 ligases for the KIH degron, we used CRISPR-Cas9 to obtain monoclonal populations of UBR4 knock-out cells (UBR4 KO, Supplemental Figure 3A). We also received monoclonal populations of UBR1 and UBR2 double knock-out cells (UBR1/2 KO) (34). We then expressed our ubiquitin-fusion constructs in parental cells, UBR1/2 KO cells and UBR4 KO cells (Figure 4B). In line with previous work, levels of *Renilla* with an N-terminal KIH increased ∼2.7-fold (*p* < 10^−3^) in UBR1/2 KO cells and in ∼1.5 fold in UBR4 KO cells respectively, indicating redundant targeting of primary lysine motif by different UBR E3 ligases.

The situation was different for reporters with TKIH or VKIH N-termini. Here, reporter levels increased ∼2.3- and ∼ 2-fold (*p* < 10^−4^ and *p* < 10^−3^, respectively) in the UBR4 KO cells, as opposed to 1.4- and 1.3-fold (*p* = 0.005 and *p* = 0.085, respectively) in the UBR1/2 KO ones. Encouragingly, stabilization of our KIH luciferase reporter only occurred when upon loss of the initiator methionine (MTKIH vs TKIH) in line with our results showing that the removal of the initiator methionine is necessary for decay.

Furthermore, a reliance on UBR4 was consistent with the degron rules we identified above. That is, luciferase versions bearing a second position threonine or valine, but not alanine or serine, were stabilized upon the loss of UBR4 (Figure 4C). Similarly, reporters with the related RIH degron at position 3–5, but not the AIH tripeptide, were also stabilized upon loss of UBR4 (Figure 4D), which is consistent with previous studies showing that UBR4 can also target position 2 arginines following a methionine (5). In line with these observations, when we expressed our single or double alanine mutation reporters in UBR4 KO cells, only reporters with the lysine at position 3 increased in expression, indicating that repression we observed for non-lysine containing reporters was mediated by a different mechanism (Supplemental Figure 3B, see KIH, KAH, KIA, and KAA). Lastly, we expressed our reporters with KIH at different positions in either parental or UBR4 KO cells, and found only KIH at positions 3-5 was stabilized in the absence of UBR4, further confirming the position dependence of the KIH degron (Supplemental Figure 3C). These results are consistent with previous studies showing that UBR4 plays a more prominent role than UBR1 and UBR2 in targeting position 2 lysine and arginine residues (5).

As a complementary experiment, we tested the ability of UBR4 to regulate the KIH degron in other cell types. We used the CRISPRi repression system in K562 cells to stably knockdown UBR4 (Supplemental Figure 3D). We then expressed our WT and KIH luciferases in these cells and compared expression between cells with non-targeting guides or UBR4 guides. As expected, knockdown of UBR4 in K562 cells stabilized KIH luciferase levels (Figure 4E). Thus, we conclude that UBR4 acts the primary N-recognin for the KIH degron.

Recent work has shown that UBR4 is in a complex with KCMF1 (24, 25). We therefore wondered if the UBR4-KCMF1 complex was necessary to degrade our KIH luciferase, or if UBR4 alone was sufficient. To this end, we depleted KCMF1 using CRISPRi (Supplemental Figure 3C) and compared luciferase expression between NT cells and KCMF1 KD cells (Figure 4F). As with the UBR4 depletion, loss of KCMF1 stabilized KIH luciferase levels (*p* < 10^−3^; Figure 4F); suggesting that these degrons are likely recognized and controlled by the UBR4-KCMF1 complex.

Taken together, we conclude that the KIH family of degrons, when following a position 2 threonine or valine, function similarly to known primary type-1 N-degrons and are recognized by UBR4 following methionine removal. To our knowledge, this is the very first example of a primary Arg/N-degron pathway that is exclusively mediated by methionine removal by MetAP2.

### Position 3 lysines and arginines can broadly function as N-terminal degrons in endogenous genes

Thus far, our results have shown that the KIH degron has the capacity to regulate exogenous reporters, but do little to speak to the role of this degron for endogenous genes. To explore this idea, we made use of a previously published dataset that quantified the impact of endogenous N-termini on protein stability (5). Here, Timms and colleagues again used the ubiquitin-fusion system and determined a “protein stability index” for each N-terminus with or without the position 1 methionine. Consistent with their observations, re-analysis of their dataset showed that N-termini containing MK were unstable (Figure 5A). Strikingly, there was no significant difference in PSI between MK and TK (or VK) N-termini (Mann-Whitney p-value = 0.953 and 0.112, respectively), indicating that these N-terminal motifs are broadly destabilizing across the proteome. In contrast and consistent with our analyses (Figure 3), AK and SK N-termini were significantly stabilized compared to MK termini, (Mann-Whitney p-value = 0.03, and *p* < 10^13^ respectively), and we observed a similar trend with arginine termini, MR, TR, VR, AR, and SR (Supplemental Figure 4A). Moreover, given the importance of methionine removal for these degrons, we calculated the difference in PSI values for N-terminal fusions with methionine from the same N-terminal fusions that lacked methionine (where a higher value indicates more instability for the no-methionine version). We then sub-grouped N-termini into those with a position 3 lysine or any other amino acid (excluding arginine). N-termini bearing MT or MV showed a greater loss of stability upon methionine removal with a position 3 lysine (*p* < 10^−3^ and *p* < 10^−4^ respectively, Figure 5B); but this difference was not seen for N-termini with alanine or serine at position 2, exactly in line with what we observed with our luciferase reporters. We performed the same analysis on position 3 arginines, and found a similar trend (Supplemental Figure 4B). Thus, these data suggest that the MetAP2–UBR4 pathway for position 3 lysines and arginines can function on endogenous N-termini.

**Figure 5.**
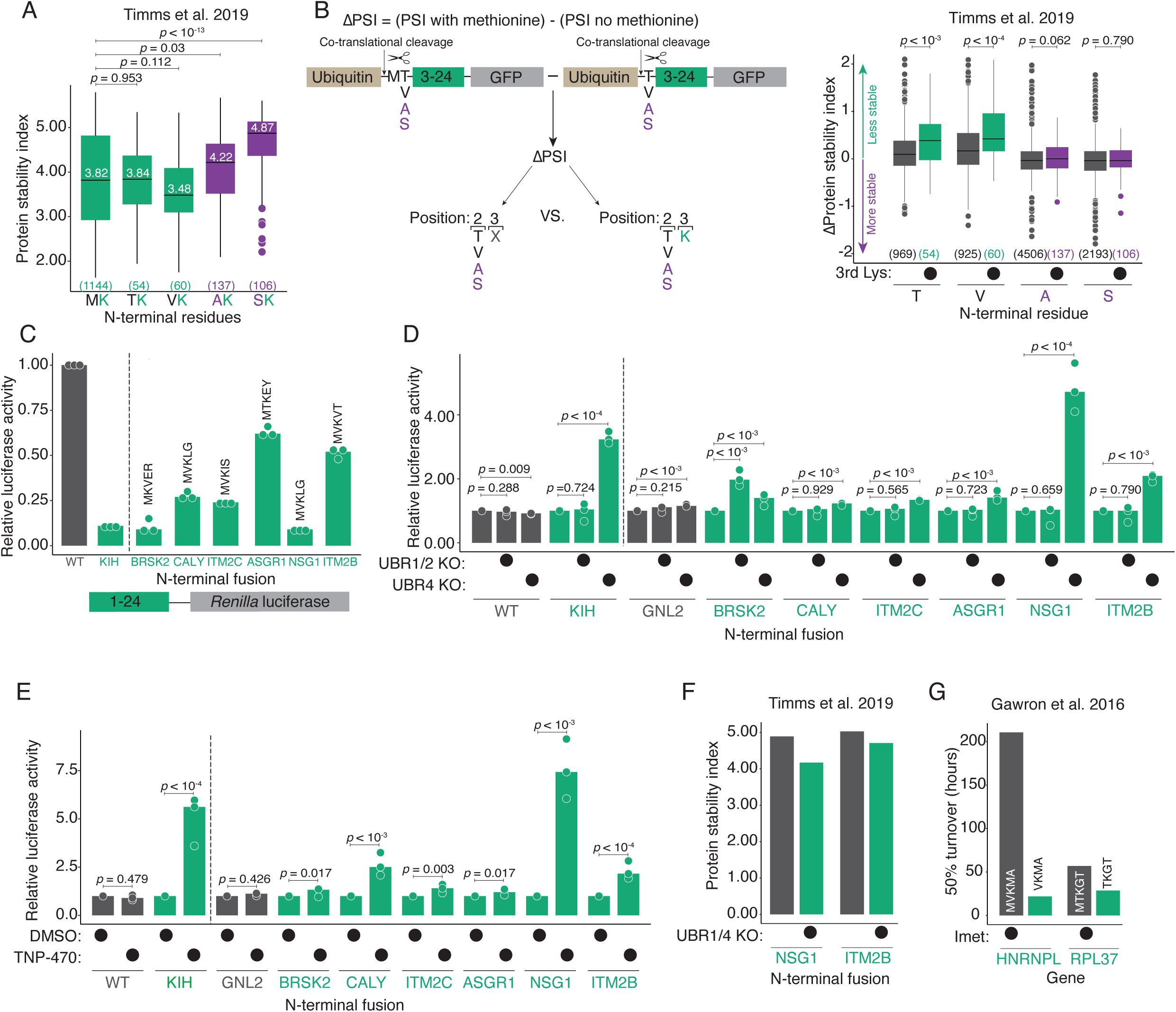
Position 3 lysines can broadly function as N-terminal degrons in endogenous genes. (A) Position 3 Lys following position 2 Thr and Val destabilize endogenous proteins. Boxplots comparing the distribution of stabilities of MK, TK, VK, AK and SK termini from Timms *et al.* 2019. The median stability for each N-termini is displayed in white, and the number of observations for each N-termini is displayed below each boxplot in parentheses. P-value was calculated using the Mann-Whitney U test. (B) Loss of N-terminal methionine destabilizes endogenous proteins that contain a position 3 Lys following a position 2 Thr or Val. Schematic of the analysis shown to the left of the plot. The N-terminome GPS data was filtered for proteins that contained either a position 2 Thr, Val, Ala or Ser. Proteins were then sub-grouped into those that contained a position 3 Lys or any other amino acid at position 3 excluding Arg. ΔPSI was calculated by subtracting the PSI of the protein when it lacked methionine from the PSI of the same protein when it contained methionine. Boxplots compare the distribution of ΔPSI values for each position 2 N-terminal amino acid. The number observations for each N-termini are plotted below the respective boxplot, and P-value was calculated using the Mann-Whitney U test. (C) K3 reporters display a range of expression levels. Schematic of the constructs shown below the plot. The first twenty-four amino acids of the indicated endogenous proteins were fused to a small flexible linker, and then to a WT Renilla luciferase reporter that contained a minimal EMCV IRES and firefly luciferase. Reporters were transfected into parental HEK293T cells, and *Renilla* luciferase activity was determined relative to the firefly control. *Renilla* expression was calculated relative to the WT *Renilla* luciferase variant. The bar represents the median activity, and each dot represents one biological replicate (composed of three technical replicates). P-value was calculated using the Mann-Whitney U test. (D) UBR4 regulates the expression of NSG1 and ITM2B K3 reporters. As in (C) except the reporters were transfected into parental, UBR1/2 or UBR4 KO cells, and *Renilla* expression for each reporter was calculated relative to the expression of the same reporter in parental cells. (E) MetAP2 regulates the expression of CALY, NSG1, ITM2B K3 reporters. As in (D) except, reporters were only transfected into parental cells. Twenty fours after the addition of the inhibitor, relative luciferase activity was quantified as in (A) and *Renilla* luciferase expression was normalized to DMSO treated cells. (F) Removal of the N-terminal methionine is necessary for UBR4 to target NSG1 and ITM2B K3 reporters. Analysis of PSI of GPS NSG1 and ITM2B reporters from Timms *et al.* 2019 when expressed in NT or UBR1/4 KO cells. Raw PSI values are plotted for each construct. (G) Removal of the initiator methionine decreases the 50% turnover rate of endogenous proteins with a position 3 Lys. Comparison of the 50% turnover rate for HNRNPL and RPL37 proteins with or without methionine from Gawron *et al*. 2016.

To further explore this possibility, we selected five N-termini to test directly using a series of criteria: first, the N-termini had to contain either MTK or MVK as the primary isoform defined by UNIPROT; second, they had to have been tested in the previous N-terminome study; and third, we selected five N-termini with greatest loss in stabilization upon methionine removal. These proteins were CALY, ITM2C, ASGR1, NSG1 and IT2MB. As a positive control for targeting by UBR E3 ligases, we selected BRSK2 isoform 6 (N-terminus: MKVER) which is known to be stabilized in the absence of UBR1 and UBR2 (5). As a negative control for regulation by MetAP2, we selected GNL2 (N-terminus: MVKPK) which showed no difference in PSI upon methionine removal (5). We then constructed a set of luciferase reporters where we fused the first 24 amino acids of these proteins to a short linker (ASTAGLT) and then *Renilla* luciferase, followed by EMCV IRES and firefly luciferase as an internal control. One key difference between these reporters and the previously published ones is that ours lacked an upstream ubiquitin and enabled us to directly interrogate the role of methionine removal by MetAP2 (5).

We transfected these N-terminal fusions along with our original wild-type and KIH variants into parental, UBR1/2 KO and UBR4 KO cells. First, we examined the expression of N-terminal fusions relative to the wild-type *Renilla* in parental cells (Figure 5C). We observed a range of luciferase levels for the N-terminal fusions, with the BRSK2 fusion being the most lowly expressed. Moreover, we found that the *Renilla* luciferase levels of the N-terminal fusions almost perfectly correlated with the known half-lives of the entire proteins (Pearson’s correlation *R* = 0.99, *p* = 0.008, Supplemental Figure 4C), consistent with these effects being due to the action of the N-termini on the stability of the *Renilla* reporter.

We then compared the expression of the reporters across cell types, normalizing expression to parental cells (Figure 5D). Consistent with our expectations, we observed the BRSK2 variant was significantly stabilized in the UBR1/2 KO cells (median fold-change = 1.97, *p* < 10^−3^). On the other hand, loss of UBR4 reliably stabilized the KIH reporter, along with BRSK2 (median fold-change 1.40, *p* < 10^−3^), and two K3 N-terminal fusions NSG1 (N-terminus: MVKLG, median fold-change = 4.72, *p* < 10^−4^) and ITM2B (N-terminus: MVKVT, median fold-change = 2.09, *p* < 10^−3^). To test the role of methionine removal by MetAP2, we expressed our reporters in parental cells and inhibited MetAP2 with TNP-470 as before (Figure 5E). As expected, MetAP2 inhibition significantly increased the levels of NSG1 and ITM2B reporters (median fold-change = 6.7 and 2.2, respectively, *p* < 10^−4^). Consistent with these results, upon re-analysis of data from Timms and colleagues, we found that both NSG1 and ITM2B N-terminal fusions with an intact position 1 methionine showed minimal differences in stability between cells that were transduced with a non-targeting guide and cells that lacked both UBR1 and UBR4 (UBR1/4 KO; Figure 5F). Thus, we conclude that these fusion proteins undergo the same mechanism of decay as KIH Renilla luciferase, which is initiated by methionine removal and leads to recognition of the lysine degron by UBR4.

Nonetheless, we were surprised to see that only two of five tested K3 reporters were stabilized upon UBR4 depletion, and we wondered if the N-terminal methionine might not be cleaved from some of our N-terminal fusions, thereby preventing recognition by UBR4. Consistent with this possibility, two of the fusions (ITM2C and ASGR1) showed no stabilization upon MetAP2 inhibition. On the other hand, one N-terminal fusion, CALY (N-terminus: MVKLG) that was not stabilized in our UBR4 KO cells was stabilized upon MetAP2 inhibition (median-fold change = 2.5, *p* < 10^−3^) despite having the same N-terminal pentapeptide sequence as NSG1. Thus, methionine removal from MTK or MVK peptides is necessary but not sufficient to trigger decay by this pathway, and downstream amino acids play a further role in E3 ligase specificity. Additionally, we performed the same experiments using position 3 arginine (R3) N-terminal fusions that fit the same criteria highlighted above (Supplemental Figure 4D). As expected, our positive control, RAD51C (N-terminus: MRGKT) was stabilized (1.5-fold, *p* < 10^−3^) upon the loss of UBR1 and UBR2 as has been shown previously (5). Interestingly, one R3 fusion, ENOSF1 (N-terminus: MVRGR) was also slightly stabilized in UBR1/2 KO cells (median fold-change = 1.30, *p* = 0.002) but not in the UBR4 KO cells. On the other hand, we observed stabilization of ZCCHC9 (N-terminus: MTRWA, median fold-change = 1.40, *p* < 10^−3^) in the UBR4-depleted cells but not cells that lacked UBR1 or UBR2.

Lastly, we searched for examples of entire proteins whose expression might be differentially regulated by this pathway. To this end, we re-analyzed steady-state mass spectrometry data of parental vs UBR4 KO HEK293T cells (Supplemental Figure 5A)(35). Here, we found a single protein, DNAJA1 (N-terminus: MVKET), that contained the K3 Arg/N degron motif and whose expression was upregulated upon loss of UBR4. Additionally, we turned to other studies where Gawron and colleagues measured the 50% turnover rates of N-terminal proteoforms upon methionine processing (36). We searched their data for proteins whose N-termini consisted of a threonine or valine at position 2 or a lysine or arginine at position 3, as well as proteins that had an observed turnover rate for both the methionine containing and methionine cleaved proteoforms. Only two proteins in their dataset fit these criteria: HNRNPL (N-terminus: MVKMA) and RPL37 (N-terminus: MTKGT). Consistent with our model, loss of methionine in both proteins strongly decreased the 50% turnover rates of these proteins (Figure 5G). Taken all together, these results show that there are endogenous N-terminal sequences whose degradation is mediated by a MetAP2-dependent UBR4 Arg/N-degron pathway.

## DISCUSSION

To uncover the role of N-terminal amino residues in controlling gene expression, we inserted KIH into positions 3-5 of reporter constructs. Unexpectedly, based on the then-conception of N-degrons, the insertion of these residues triggered the rapid decay of *Renilla* luciferase by the ubiquitin proteasome system. A series of rational mutations at each position revealed that this decay was mediated by the properties of the amino acids: a positive charge at position 3, semi-bulky hydrophobic residues at position 4, and aromatic or bulky residues at position 5 conferred the strongest loss of expression. This degron family represents a newly described Arg/N-degron pathway, whereby MetAP2 removes the initiator methionine of MTK/R and MVK/R N-terminal peptides, thus shifting the position of the lysine/arginine from 3 to 2 and making that residue accessible for recognition by UBR4-KCMF1. Finally, our analysis of endogenous N-termini indicates that this pathway is not confined to N-terminus of *Renilla* reporter. Thus, our results add an additional layer of regulation to existing Arg/N-degron pathways, showing how position 3 residues can become targets of UBR box E3 ligases upon methionine processing.

One surprise of our results was the differential impact of MetAP1 and MetAP2 cleavage on this degron family. That is, the removal of the N-terminal methionine, in and of itself, was not sufficient to induce the decay of peptides that have a position 3 N-terminal lysine or arginine. For example, we found the KIH degron functioned robustly when preceded by threonine or valine but not alanine or serine. While we did not resolve these differences, one possibility is that subsequent N-terminal acetylation of the newly exposed alanine and serine termini prevents recognition by UBR4-KCMF1. In support of such a possibility, global studies of N-terminal acetylation show that 95% and 99% of alanine and serine N-termini are co-translationally acetylated, whereas N-terminal acetylation occurs for only 82% and 20% of threonine and valine N-termini (37). Recent work has also shown that acetylation of the N-terminal methionine residues prevented the recognition of type-2 N-degrons by UBR4-KCMF1 (11). Of course, we cannot formally rule out the possibility that there are some AK and SK termini that are degraded in a similar manner, and the interplay between second-position preferences for MetAP1, MetAP2, and acetyltransferases highlights the complexity of such “rules” for N-degrons.

This family of N-degrons appears to be primarily recognized by the UBR4 E3 ligase rather than UBR1 and UBR2. This finding is in line with structural differences in the UBR box domains as well as the broader substrate selectivity for position 2 K and R residues (MK and MR) by UBR4 than UBR1 and UBR2 (5, 21, 22). As with other aspects of UBR4 targeting including type-2 N-degrons and internal degrons (2, 5, 11, 19, 21, 24), KIH and related motifs require the UBR4-KCMF1 complex for degradation. Interestingly, these results help highlight a divergence in degrons and E3 ligase selectivity within eukarya. That is, in yeast, reporters bearing MK, TK and VK N-termini were not unstable nor were they further stabilized in the absence of Ubr1 (7). Unlike humans, yeast only has one UBR E3 ligase (Ubr1), which is structurally most similar to human UBR1 and UBR2, not human UBR4 (21); therefore, it is tempting to speculate that the duplication of UBR E3 ligases and their structural differences are important contributors to the possible expansion of N-degrons.

Lastly, we discovered that endogenous N-termini are subject to this mechanism of decay. Analysis of N-terminal proteoforms support our mechanism, as we observed a handful of peptides whose 50% turnover time markedly decreases when methionine is removed exposing N-terminal TK and VK dipeptides. Additionally, steady-state proteomics comparing WT to UBR4 KO cells highlights additional targets that may be substrates of the MetAP2-dependent Arg/N-degron pathway. Nonetheless, our analysis of endogenous N-termini further underscores the complexity of these pathways. Of the five of N-terminal fusions we tested, two were degraded as expected, two did not undergo methionine removal by MetAP2, and one (CALY) was degraded by unknown mechanisms. In other words, downstream amino acids (e.g., in ITM2C and ASRG1) also appear to play a role in methionine removal through an unknown mechanism, in addition to impacting E3 ligase specificity.

As has been known, the logic of N-degron pathways is highly complex, and our results highlight the complicated dynamics between MetAP1 and MetAP2 specificity, the role and influence of N-acetylation, redundancy, and divergence of UBR E3 ligases, and the availability of ubiquitin-accepting lysine. As a field, we will need to integrate multiple sequence determinants to be able to predict the impact of N-termini on protein decay, and we hope that our results motivate further unbiased studies, especially with state-of-the-art large language models. Nonetheless, in the long term, such rules will need to be further combined with studies of translation, including start site selection and clearance, to reach a full understanding of how this important, dual-purpose region regulates gene expression.

## MATERIALS & METHODS

### Lead contact

Further information and requests for resources and reagents should be directed to and will be fulfilled by the lead contact, Olivia Rissland.

#### Mammalian cell culture

HEK293T cells and U2OS cells were grown and maintained in Dulbecco’s modified eagle medium (DMEM) with 25 mM D-glucose, 4 mM L-glutamine, 3.7 g/mL NaHCO_3_, 110 mg/mL sodium pyruvate, and supplemented with 10% v/v standard fetal bovine serum (FBS, Gibco). Zim3-dCas9 K562 cells (a kind a gift from Marco Jost’s laboratory, Harvard University) were grown and maintained in RPMI 1640 medium with 2 mM L-glutamine (Gibco) and supplemented with 10% v/v standard FBS (Gibco), 25 mM HEPES (Gibco), and 2 mM GlutaMAX (Gibco). Zim3-dCas9 RPE1 cells (a kind a gift from Marco Jost’s laboratory, Harvard University) were grown and maintained in DMEM: F12 (Gibco) supplemented with 10% v/v standard FBS (Gibco), and 0.01 mg/mL hygromycin B (ThermoFisher). CHO-K1 cells (ATCC-CCL61) were grown and maintained in Kaighn’s modification of Ham’s F-12 medium (F-12K medium, ATCC) with 2 mM L-glutamine, 1.5 g/mL NaHCO_3_, and supplemented with 10% v/v standard FBS (Gibco). NIH3T3 cells (ATCC, CRL-1658) were grown and maintained in DMEM with 25 mM D-glucose, 4 mM L-glutamine, 3.7 g/mL NaHCO_3_, 110 mg/mL sodium pyruvate, and supplemented with 10% v/v standard donor bovine serum (DBS, Gibco). All cells were grown at 37 °C in a 5% CO_2_ humidified incubator.

### Transfections

HEK293T, U2OS, CHO-K1, and NIH3T3 cells were transfected with Lipofectamine 2000 (ThermoFisher) according to manufacturer’s instructions at a ratio of 1:5 of DNA (in μg) to Lipofectamine 2000 (in μL). RPE1 cells were transfected with Lipofectamine 3000 (ThermoFisher) at a ratio of 1:5 of DNA (in μg) to Lipofectamine 3000 (in μL), and a ratio of 1:10 of DNA (in μg) to PLUS reagent (in μL). K562 cells were reverse transfected with Lipofectamine LTX (ThermoFisher) according to manufacturer’s instructions at a ratio of 1:5 of DNA (in μg) to Lipofectamine LTX (in μL), and a ratio of 1:10 of DNA (in μg) to PLUS reagent (in μL). Cells were harvested 24 hours after transfection unless stated otherwise.

### Lentivirus production

Lentivirus was generated by transiently transfecting ∼7,000,000 HEK293T cells with 6 μg of transfer plasmid, 4.5 μg of psPAX2 (a kind gift from Mikko Taipale), and 750 ng of VSV-G (a kind gift from Mikko Taipale) with 30 μL of Lipofectamine 2000 transfection reagent. Twenty-four hours after transfection, the medium was aspirated and replaced with fresh medium. Viral supernatant was then collected 48 hours later by filtration through a 0.45 μm PES filter (Millipore Sigma) and stored at –80 °C prior to transduction.

### Cell line generation

UBR1/UBR2 knockout (UBR1/2 KO) HEK293T cells were a kind gift from Alexander Varshavsky and Marcia Goldberg, and were generated as previously described (34).

UBR4 knockout HEK293T cells (UBR4 KO) were generated similarly to UBR1/2 KO cells. Briefly, HEK293T cells were transiently transfected with 1 μg of the lentiCRISPR v2 (Addgene #52961) plasmid containing a previously validated UBR4 sgRNA or a non-targeting sgRNA (5). Transformants were selected with 2 μg/mL puromycin for 48 hours, and individual clones were expanded with limiting dilution (0.5 cells/well) in 96-well plates. Loss of gene expression was verified by immunoblotting.

UBR4 sgRNA: GCCTCTCGAAGATGAACACCG
NT sgRNA: ATCGTTTCCGCTTAACGGCG

Zim3-dCas9 K562 cells were a kind gift from Marco Jost and generated as described previously (33). To generate UBR4, MetAP2, and KCMF1 knockdown lines, cells were transduced at an MOI < 0.3 with virus made via the pJR103 dual guide RNA plasmid (a kind gift from Marco Jost & Jonathan Weissman, Addgene #187242) (33). Each plasmid contained 2 unique gRNAs targeting the gene of interest that were empirically validated in previous CRISPRi screens or non-target (NT) guides (33).

sg-NT1: GGAGTTAAGGCCTCGTCTAG
sg-NT2: GTCCCAGGCTCTCCACTATG
sg-NT3: GGCGGGCGCAAGACGTGGCA
sg-NT4: GGCTAGAACCCACACTCTTA
sg1-UBR4: GCGTTGGCGGAGGGAAACCC
sg2-UBR4: GCTACTGCGGCTCCCTCCGG
sg1-MetAP2: GCCTCCGGGAGCCACCTGAA
sg2-MetAP2: GTCGGGCCCAGCGACCCCAG
sg1-KCMF1: GCAGCGCCGGGACCCCGCGG
sg2-KCMF2: GGACATCCTAGTTCAGACGG

#### Plasmids

All plasmids used in this study are listed in Supplemental table 1. A detailed description of plasmid construction is contained in an additional supplemental file.

#### Immunoblotting

Cells were washed gently with ice-cold phosphate buffered saline (PBS), harvested into 1.5 mL microcentrifuge tubes, and then pelleted by centrifugation at 500*g* for 5 minutes. The cells were then lysed with lysis buffer A (100 mM KCl, 0.1 mM EDTA, 20 mM HEPES, pH 7.6, 0.4% NP-40, and 10% glycerol v/v) supplemented with 1 mM DTT and 1 x cOmplete EDTA-free protease inhibitor cocktail (Roche). Lysates were then clarified by centrifugation at 14000*g* for 5 minutes. Total protein concentration was calculated using Pierce™ Detergent Compatible Bradford Assay Kit (ThermoFisher) according to manufacturer’s instructions. Lysates were then prepared with 4x NuPAGE LDS Sample Buffer (Invitrogen) and 10x Bolt Sample Reducing Agent (Invitrogen) and boiled at 95 °C for 5 minutes. 20-30 μg of protein from cell lysates was resolved on NuPAGE 4-12%, Bis-Tris, 1.0 mM, Mini Protein precast gels (Invitrogen) with 1x MES SDS Running Buffer (Invitrogen) at 160 V for 50 minutes at room temperature. Protein was transferred to a PVDF membrane (Amersham), according to the manufacturer’s instructions at 20 V for 60 minutes. For UBR4 immunoblots, 20-30 μg of protein from cell lysates was resolved on NuPAGE 3-8%, Tris-Acetate, 1.0-1.5 mm, Mini Protein precast gels (Invitrogen) at 160 V for 50 minutes. Protein was then transferred to a PVDF membrane (Amersham), according to manufacturer’s instructions at 20 V for 12-16 hours at 4**°**C.

The membrane was then blocked with TBST (1x TBS with 0.1% v/v Tween) with 3% w/v nonfat powdered milk and rocked at room temperature for 30 minutes. Antibodies were diluted in TBST with 3% w/v nonfat powdered milk, and blots were incubated by rocking overnight at 4 **°**C. The following day, the membranes were washed three times with TBST and then incubated with the corresponding horseradish peroxidase (HRP) linked secondary antibody for 1 hour at room temperature. Membranes were then washed three times with TBST, and protein signal was detected with ECL (Cytiva) and imaged on Sapphire Bimolecular Imager (Azure). Western intensity was quantified using Fiji ImageJ.

Antibodies were diluted as follows: anti-UBR4 (anti-p600, Fortis Life Science), 1:1000, anti-UBR1 (Santa Cruz Biotechnology) 1:1000, anti-UBR2 (Fortis Life Science) 1:000, anti-Renilla luciferase 1:500 (Novus Biologicals), anti-GFP (Sigma Aldrich) 1:2000, anti-Beta actin 1:2000 (Cell Signaling).

#### Luciferase assays

Luciferase assays were performed using Dual-Luciferase Reporter Assays System (Promega) according to manufacturer’s instructions. Briefly, after experimentation cells were removed from the incubator, the media was aspirated, and the cells were stored at −80**°**C for at least 24 hours. Cells were suspended in 1x passive lysis buffer and rocked at room temperature for 15 minutes. After lysis, 20 μL of lysate was loaded into a white 96-well flat bottom plate (Genesee Scientific). Luciferase activity was then measured with a GloMax Navigator. For each experiment, three biological replicates, each with three technical replicates (three different wells transfected on the same day) were performed. Relative luciferase activity was then quantified by taking the ratio of *Renilla* luciferase to firefly luciferase.

#### RT-qPCR

Cells were washed gently with ice cold PBS, harvested into 1.5 mL centrifuge tubes, and pelleted at 500*g* for 5 minutes. Cells were then re-washed with PBS and then suspended in 1 mL of TRIzol Reagent (Invitrogen). RNA was then extracted according to manufacturer’s instructions. RNA (2 μg) was treated with TURBO DNase (ThermoFisher) for 1 hour, and then reverse transcribed with SuperScript IV Reverse Transcriptase and 2μM of Random Hexamers (ThermoFisher) according to manufacturer’s instructions. RT-qPCR was performed with iTaq Universal SYBR Green Supermix (Bio-Rad) using the LightCycler 480 (Roche).

#### *In vitro* translation

*In vitro* translation assays were performed in nuclease-treated rabbit reticulocyte lysate (Promega). Reporters containing either the wild-type or the KIH *Renilla* luciferase variant flanked by an EMCV IRES and a firefly luciferase, were amplified by PCR to include the T7 promoter. PCR products (200 ng) were then *in vitro* transcribed with the mMESSAGE mMACHINE T7 Transcription Kit (ThermoFisher) according to manufacturer’s instructions. Samples were then treated with 1 μL of TURBO DNase (ThermoFisher) for 30 minutes at 37**°**C. Messenger RNA was then polyadenylated with Poly(A) Tailing Kit (ThermoFisher) according to manufacturer’s instructions. The polyadenylated mRNA was purified by LiCl (ThermoFisher, 2.5 M final concentration) precipitation according to manufacturer’s instructions, and incubated in a 20 μL rabbit reticulocyte lysate mixture that contained: 1 μg of mRNA, 14 μL of rabbit reticulocyte lysate, 10 mM Amino Acid Mixture minus Methionine, 10 mM Amino acid Mixture minus Leucine and nuclease free water. The reaction was incubated at 30**°**C for 90 minutes and then diluted 1:10 using nuclease free water. Luciferase expression levels were then measured using the dual luciferase assay, and platting each sample in technical triplicates.

#### Inhibitors

Cells were treated with inhibitors for 24 hours unless stated otherwise. For MG132 (Millipore Sigma) experiments, MG132 was used at a final concentration of 10 μM. For cycloheximide chase experiments, cycloheximide (ThermoFisher) was used at a final concentration of 80 μg/mL. For MetAP2 inhibition experiments, TNP-470 (Millipore Sigma) was used at a concentration of 50 nM.

## BIOINFORMATICS

### HEK293T endogenous short half-life analysis

Analysis of short-lived proteins in HEK293T cells from Li et al. 2021 (Table S2 in their paper) was performed with R (28). To create a density plot of short-lived proteins (Figure 1H), the file uploaded into R, and the data was filtered for proteins with half-lives less than 8 hours. To compare relative luciferase activity to endogenous half-lives (Supplementary Figure 4C), we filtered the data for proteins that had a measured half-life that matched our K3 reporters.

### Human N-terminal global protein stability analysis

Analysis of the stability of N-terminome via the global protein stability assay from Timms et al. 2019 (Supplementary File 1 in their paper) was performed with R (5). Briefly, the data frame was imported into R and sub-grouped into two data frames: proteins that contained N-terminal methionine, and proteins without the N-terminal methionine. The ΔPSI for a given protein was calculated by subtracting the PSI of that protein when it lacked the N-terminal methionine from the PSI of the same protein (matched by Ensembl Transcript ID) when it contained the N-terminal methionine: ΔPSI = PSI (with methionine) – PSI (no methionine). Data was then filtered to include N-termini that contained position 2 amino acids Lys, Arg, Thr, Val, Ala, and Ser. The distribution of stabilities (either PSI or ΔPSI) or was then plotted for each N-termini of interest.

### Turnover rate of endogenous proteins analysis

Analysis of the turnover rate of proteoforms in Jurkat cells (Table EV1B in their paper) was performed in R (36). The data set was loaded into R and filtered for proteins that contained a position 2 Thr, Val, Ala or Ser, followed by a position 3 Lys or Arg and that had a measured differential turnover rate upon loss of methionine. The 50% turnover time of each proteoform was plotted.

### Mass spec analysis

Analysis of parental vs UBR4 KO HEK293T cells was performed with the Differential Enrichment analysis of Proteomics data (DEP) package in R (35, 38). The proteinGroups file was imported into R for differential enrichment analysis and the UBR4 KO1 sample was excluded due to UBR4 contamination (35). The raw dataset (5396 proteins) was filtered for contaminant proteins, decoy database hits, reverse, and proteins only identified by site. We then filtered for proteins that were quantified in two out of three replicates of at least one condition. The data was then normalized with a variance stabilizing transformation. Missing data was then imputed by assuming a normal distribution with a downshift of 1.8 standard deviations with of 0.3 the original normal distribution. To determine differentially expressed proteins, false discovery rate was set to 0.05 and a fold-change of 1.5.

#### Data visualization, quantification, and statistical analysis

Data visualization, quantification, and statistical analyses were conducted in R. Visualization was performed using the ‘ggpubr’ package and plots were further refined in Adobe Illustrator (39). Statistics were performed using the ‘rstatix’ package (40).

## ACKNOWLEDGEMENTS

We would like to thank Eric Bennett, Sujatha Jagannathan, Michael Rapé, Sichen (Susan) Shao, and other members of the Rissland Lab for thoughtful comments and helpful discussions. We would like to thank the labs of Thomas Arnesen, Stephen Elledge, Marcia Goldberg, Marco Jost, Mikko Taipale, and Alexander Varshavsky for reagents and cell lines. This work was supported by NIH grant R35GM128680 (OSR) and the RNA Bioscience Initiative, University of Colorado School of Medicine (OSR), NIH training grant T32-GM136444 (EJM), NSF-GRFP 1000300571 (EJM) and HHMI Gilliam Award GT14919 (EJM).

## AUTHOR CONTRIBUTIONS

Conceptualization: EJM and OSR.; methodology: EJM and OSR.; investigation: EJM and EH.; writing - original draft: EJM; writing – review & editing: EJM, EH, OSR.; software: EJM; formal analysis: EJM; visualization: EJM and OSR; supervision: OSR; funding: EJM and OSR.

## SUPPLEMENTAL TABLE LEGENDS

**Supplemental Table 1. List of plasmids used in each figure.**

**Supplemental Table 2. Expanded list of PCR primers, Oligonucleotides and CRISPR guides used for cloning.**

## SUPPLEMENTAL FIGURE LEGENDS

**Supplemental Figure 1. The KIH motif is degron across mammalian cell types.** (A) The KIH motif is decreases expression regardless of codon content. HEK293T cells were transfected with either wild-type, KIH (codons: AAG ATC CAC) or KIH (codons: AAA ATA CAT) *Renilla* luciferase variants, and co-transfected with firefly luciferase as a control. *Renilla* luciferase activity was quantified relative to the firefly control, and *Renilla* expression was calculated relative to the wild-type version. The bar represents the median activity, and each dot represents one biological replicate (composed of three technical replicates). P-value was calculated using the Mann-Whitney U test. (B) Same as in A, using the KIH-AAG ATC CAC except transfection into the indicated human cell lines. (C) Same as in (B) except transfection into the non-human indicated cell lines.

**Supplemental Figure 2. CRISPRi depletion of MetAP2 in K562 cells.** (A) CRISPRi knockdown of MetAP2. Zim3-dCas9 expressing K562 cells were either untransduced, or transduced with dual non-targeting (NT) gRNAs or dual MetAP2 gRNAs. Lysates were made and run by gel electrophoresis, and membranes probed for MetAP2, and beta-actin.

**Supplemental Figure 3. UBR4 only targets position 3 lysine motifs** (A) CRISPR-Cas9 knockout of UBR1, UBR2, and UBR4 N-recognins. UBR1/2 KO cells were generated as previously described by Vu and Varshavsky 2020. HEK293T cells were transfected with either a non-targeting guide or a previously designed gRNA that targets UBR4. Transformants were selected with 2 ug/mL puromycin, and monoclonal populations were expanded. Lysates were prepared and run by gel electrophoresis, and membranes probed for UBR1, UBR2, UBR4, and beta-actin. (B) Loss of UBR4 stabilizes KIH at position 3-5. Parental or UBR4 KO HEK293T cells were transfected with either WT or KIH 3-5, 4-6, 5-7, and 7-9 luciferase variants. *Renilla* luciferase activity was quantified relative to the firefly control, and *Renilla* expression for each luciferase variant was calculated relative to the expression of that variant in parental cells. The bar represents the median activity, and each dot represents one biological replicate (composed of three technical replicates). P-value was calculated using the Mann-Whitney U test. (C) Reporters with alanine mutations that contain Lys at position 3 are recognized by UBR4. As in (B) except with the indicated reporters. (D) CRISPRi knockdown of UBR4 and KCMF1. Zim3-dCas9 expressing K562 cells were either untransduced, or transduced with dual non-targeting (NT) gRNAs or dual gRNAs targeting UBR4 or KCMF1. Lysates were made and run by gel electrophoresis, and membranes probed for UBR4, KCMF1, and beta-actin.

**Supplemental Figure 4. Position 3 arginines broadly function as N-terminal degrons in endogenous genes** (A) Position 3 Arg following position 2 Thr and Val destabilize endogenous proteins. Boxplots comparing the distribution of stabilities of MR, TR, VR, AR, and SR termini from Timms *et al.* 2019. The median stability for each N-termini is displayed in white, and the number of observations for each N-termini is displayed below each boxplot in parentheses. P-value was calculated using the Mann-Whitney U test. (B) Loss of N-terminal methionine destabilizes endogenous proteins that contain a position 3 Arg following a position 2 Thr or Val. Schematic of the analysis shown to the left of the plot. The N-terminome GPS data was filtered for proteins that contained either a position 2 Thr, Val, Ala or Ser. Proteins were then sub-grouped into those that contained a position 3 Arg or any other amino acid at position 3 excluding Lys. ΔPSI was calculated by subtracting the PSI of the protein when it lacked methionine from the PSI of the same protein when it contained methionine. Boxplots compare the distribution of ΔPSI values for each position 2 N-terminal amino acid. The number observations for each N-termini are plotted below the respective boxplot, and P-value was calculated using the Mann-Whitney U test. (C) K3 reporter expression levels correlate well with endogenous half-lives. Scatterplot comparing the relative *Renilla* luciferase activity of WT, KIH, ITM2B, and ITM2C luciferase variants to calculated half-lives of WT and KIH variants from Figure 1F and to endogenous half-lives of ITM2B and ITM2C from Li *et al*. 2021. R is the Pearson’s correlation coefficient. (D) R3 reporters are targeted by UBR1/2 and UBR4. Parental, UBR1/2 KO, or UBR4 KO HEK293T cells were transfected with the indicated R3 reporter constructs and *Renilla* luciferase activity was determined relative to the firefly control. *Renilla* expression for each luciferase variant was calculated relative to the expression of that variant in parental cells.

**Supplemental Figure 5. Steady-state mass-spectrometry of WT v UBR4 KO HEK293T cells reveals potential endogenous targets** (A) DNAJA1 is a potential target of the UBR4-MetAP2 Arg/N-degron pathway. Volcano plot of differentially expressed proteins in parental vs UBR4 KO HEK293T cells. Data re-analyzed from Hegazi *et al*. 2022.

