## Supplementary material for "UBR4 regulates a MetAP2-dependent Arg/N-degron pathway": Supp File 1

**Plasmids**

pOR391 a codon optimized WT *Renilla* luciferase was constructed by ordering a codon optimized *Renilla* luciferase Minigene from IDT that was inserted into the pUCIDT vector. The codon optimized *Renilla* luciferase was then cloned into pIS1 vector (a kind gift from David Bartel, Addgene plasmid # 12179) in place of the non-optimal *Renilla* luciferase with Gibson assembly.

KIH pOR391 was constructed by digesting pOR391 with NheI and PspOMI. The *Renilla* luciferase sequence with the inserted KIH codons (AAG ATC CAC) was ordered as a pair of oligos from IDT that contained NheI and PspOMI cut sites. The oligos were annealed and digested, and then inserted into the pOR391 vector.

pIS0, Firefly luciferase, was a kind gift from David Bartel (Addgene plasmid # 12178) (1).

pcDNA 3.1 (+) *Renilla* luciferase-EMCV IRES-Firefly luciferase was constructed with Gibson assembly by inserting the EMCV IRES from the R5 EMCV plasmid (a kind gift from Vincent Mauro, Addgene plasmid # 51733) followed by Firefly luciferase from plasmid pIS0 downstream the multiple cloning sequence (MCS) of the pcDNA 3.1 (+) vector (2). Both WT and KIH *Renilla* luciferase variants were inserted into the MCS of this vector with restriction digestion using NheI and NotI.

*Renilla luciferase variants*

KIH *Renilla* luciferase variants (Figure 2) were made with the Q5 Site-Directed Mutagenesis Kit (NEB) using either WT or KIH *Renilla* luciferase-EMCV IRES-Firefly luciferase as a template.

*Ubiquitin fusions*

pcDNA 3.1 (+) Ubiquitin-MTSKV *Renilla* luciferase-EMCV IRES-Firefly luciferase and Ubiquitin-MTKIH *Renilla* luciferase-EMCV IRES-Firefly luciferase were constructed with Gibson assembly by inserting the ubiquitin moiety from the HA-Ubiquitin plasmid (a kind gift from Edward Yeh, Addgene plasmid #18712) immediately upstream the WT or KIH *Renilla* luciferase sequence in the pcDNA 3.1 (+) *Renilla* luciferase-minimal EMCV IRES-Firefly luciferase plasmid (3). Ubiquitin fusions: TSKV, TKIH, VKIH, MKIH, and KIH plasmids were made by mutating either MTSKV or MTKIH variants plasmid with the Q5 Site-Directed Mutagenesis Kit (NEB).

*N-terminal fusions*

N-terminal fusions were constructed with Gibson assembly. The first 24 codons for a given gene were ordered as oligos from IDT. The oligos were annealed and a short linker, ATSALGT, was added to each fusion by PCR. The PCR product was then inserted immediately upstream of the *Renilla* luciferase in the pcDNA 3.1 (+) WT *Renilla* luciferase-minimal EMCV IRES-Firefly luciferase plasmid.

*CRISPR plasmids*

LentiCRISPR v2 was a kind gift from Feng Zhang (Addgene plasmid # 52961) (4). Individual gRNAs were cloned into the vector according to the lentiCRISPRv2 and lentiGuide oligo cloning protocol.

pJR103 was a kind gift from Marco Jost & Jonathan Weissman (Addgene #187242) (5). Dual guide RNAs targeting genes of interest were cloned into the pJR103 vector according to Protocol 3 cloning of arrayed dual-sgRNAs (https://www.jostlab.org/_files/ugd/1b15a0_8eb892bae6cf4683be09e3e941abe4ac.pdf).
