## Supplementary figures and images for "UBR4 regulates a MetAP2-dependent Arg/N-degron pathway"

### Supp Figure 1

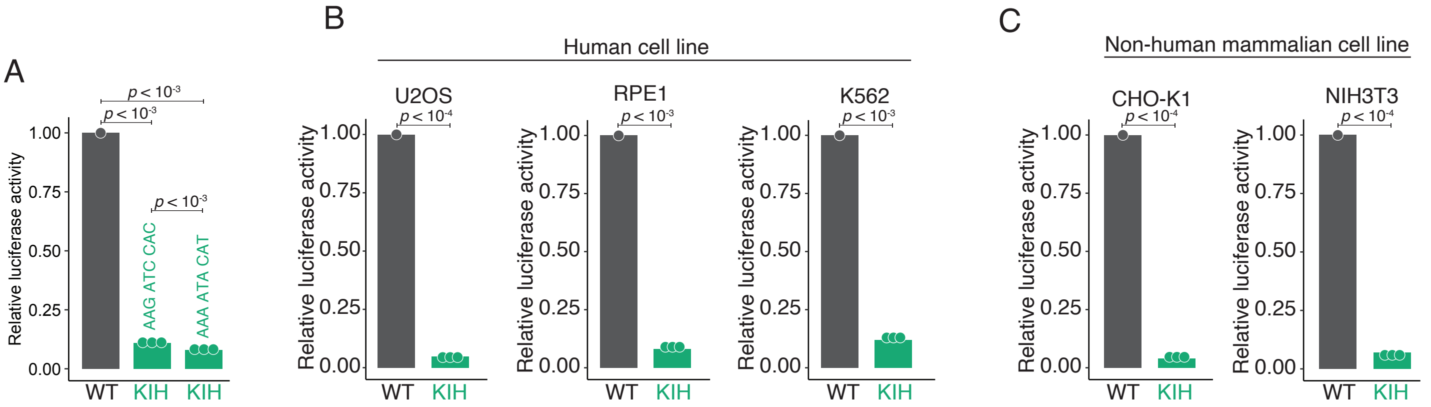

### Supp Figure 2

A

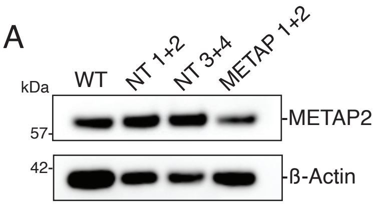

### Supp Figure 3

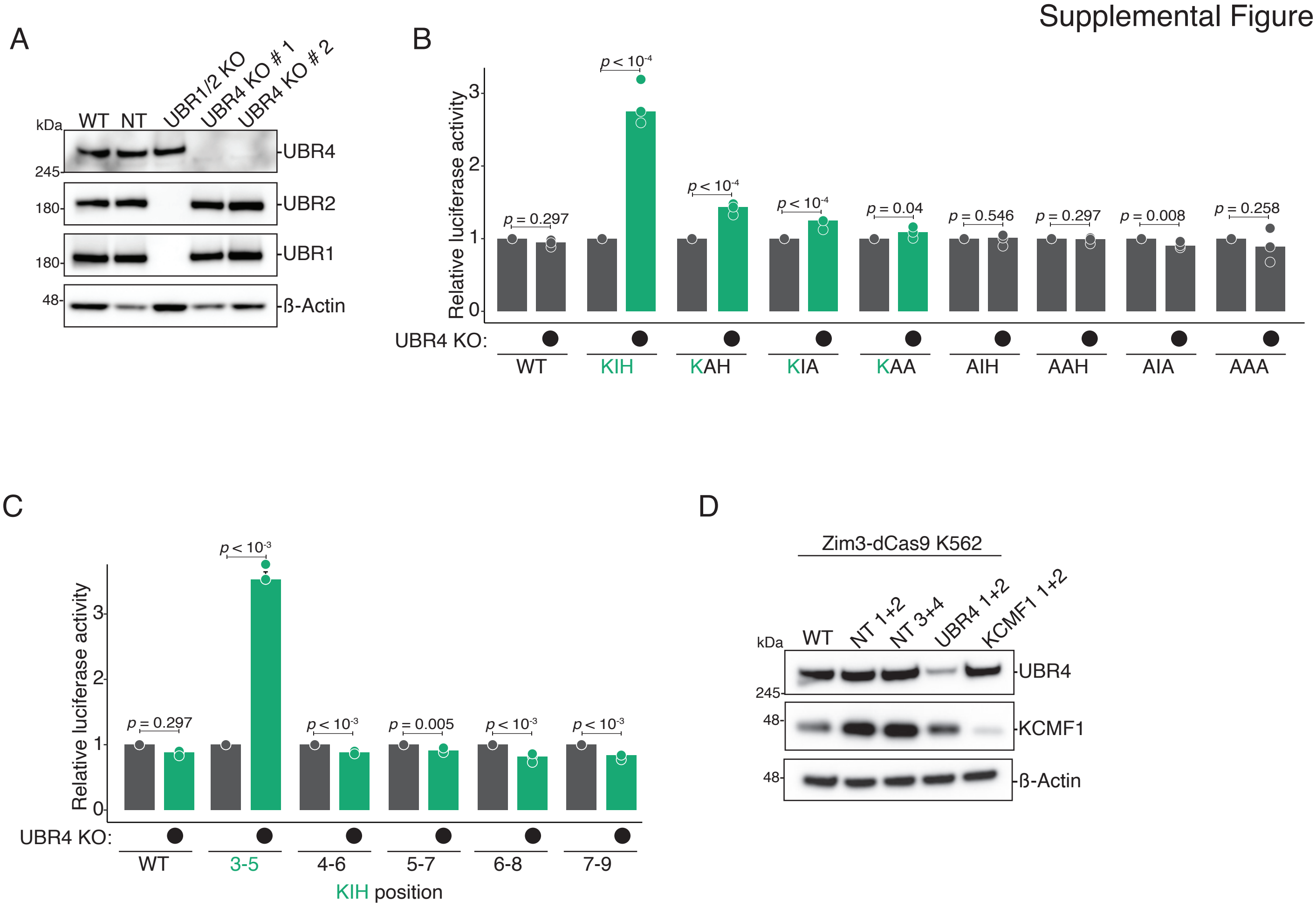

### Supp Figure 4

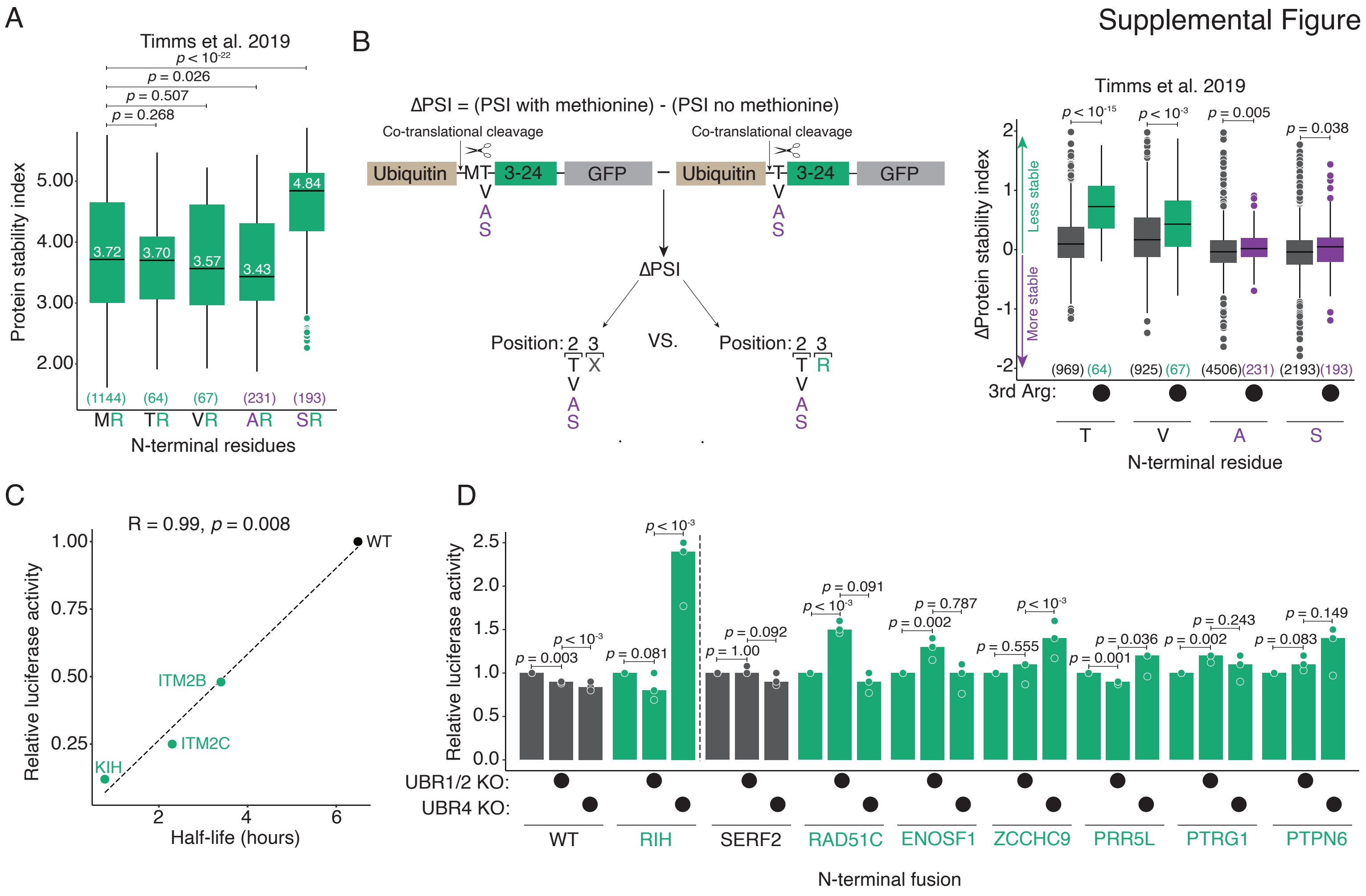

### Supp Figure 5

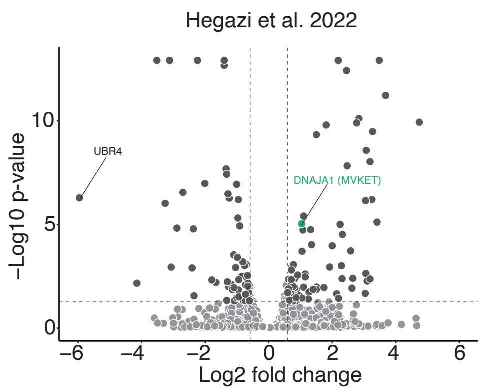
